# A New Cost Effective, Simplified Workflow for Transposon Insertion Sequencing (TIS)

**DOI:** 10.64898/2026.01.23.699905

**Authors:** Claire Hill, David Baker, John Wain

## Abstract

Transposon insertion sequencing (TIS) encompasses methods such as Transposon Directed Insertion sequencing (TraDIS) and Transposon-Sequencing (Tn-Seq); these methods are widely uses for genome scale screens of essential and conditionally essential genes. Early limitations centred on mutant generation, today the biggest factor is sequencing and data generation and the costs associated.

Here, we present and evaluate four methods TIS sequencing library generation protocols compatible with Illumina whole genome sequencing (WGS) workflows to maximise sequence efficiency and minimise turnaround times and costs. Using *E*.*coli* BW25113 transposon mutagenesis libraries generated with Tn*5* and mariner (*Himar1*) transposons, we compare the methods in terms of reagent cost, workflow complexity, and target enrichments for the recovery of unique insertion sites and essential genes identified.

All methods generated TIS data suitable for gene essentiality analysis. Illumina Flex based protocols were 4-6 tomes cheaper than the traditional Illumina Nextera based approach with similar or superior Transposon-Chromosome (Tn-Chr) junction enrichment. Across all methods, sequencing depth and mutant library density were the dominant factor in useful biological insight. Subsampling demonstrated that for a good quality mutant library, five million reads were sufficient to identify essential genes in *E*.*coli*. Whereas deeper sequencing reduced the statistical power and included contaminating background noise, seen primarily with Tn*5*.

We conclude that an Illumina Flex based approach, especially when integrating with routine WGS, provides an excellent balance of speed, cost and data quality. Assuming five million reads and a robust Illumina Flex approach, a TIS library can be sequenced for around £40.

## Introduction

Transposon Insertion Sequencing (TIS) approaches, including TraDIS (1) and Tn-Seq (2), enable high throughput, genome-wide identification of essential and conditionally essential genes. Using high throughput mutagenesis and sequencing approaches are now routinely applied to interrogate bacterial fitness under environmental and pathogenic conditions (3). When these technologies were first utilised, the cost of and throughput in mutant generation was the limiting factor but today the bottleneck is DNA sequencing for the identification of insertion sites.

The throughput of transposon mutant experiments has increased alongside the increase in sequencing capability and in some cases far more data than needed is generated. Excessive sequencing depth can bias transposon insertion counts due to PCR amplification (4), sequencing errors, or mis-mappings. This can distort the distribution of transposon insertions, making it harder to distinguish truly essential genes from those that appear essential due to technical artifacts.

Examples of TIS (rather than hybridisation) (5, 6) have been developed for most commercially available instruments: 454 pyrosequencing (7), Ion torrent (8), and Solexa (9), MinION (10) . Today, most experiments use Illumina sequencing platforms due to cost and throughout. Our experience is that the NextSeq 500 has the capacity for up to 28 TIS samples of *Escherichia coli* BW25113 mutant libraries. We now use a NextSeq 2000, producing up to 1.8 billion reads (almost 5x more data than the NextSeq 500) meaning that we could hypothetically sequence over 100 of the same libraries, using a 40% PhiX spike in to mitigate low diversity (11). There are lower output flow cells for both machines but costs per Gigabyte of data increase with the lower output; the purpose of this study is to demonstrate methods that generate TIS data with minimal costs.

This increase in capacity simplifies the need to account for variation between sequencing runs however, we rarely need to sequence 108 transposon libraries at a time. To minimise cost, and because TIS sequencing uses a unique protocol, we would batch multiple experiments; unfortunately increasing the time between experiment and results.

Furthermore, our quality control protocols for TIS include confirming the insertion density by “light touch” sequencing prior to experiments being performed; enabling optimisation of library production but delaying the generation of results even further. Thus, we have a situation where optimising TraDIS sequencing by exploiting the capacity of the sequencing machine is causing significant delays in data generation. Being able to combine TraDIS sequencing with other DNA sequencing runs would allow us to optimise for cost the sequencing service and maintain our turnaround time.

The NextSeq 2000 is routinely used for whole genome and metagenomic sequencing; here we present and evaluate a methodology for TIS sequencing that follows our minimised reaction Illumina workflow, described (12) and is therefore compatible with the preparation of libraries for standard whole genome sequencing (WGS) and amplicons. The WGS increases the diversity of the sequencing pool, allowing only 1% PhiX addition for the purpose of positive control rather than providing diversity. We present four variations on sequencing library preparation for pools of hundreds of thousands of transposon insertion mutants using two transposons (Tn*5* and mariner) and evaluate them in terms of, hands-on time in the laboratory, cost, target sequencing and genome mapping.

## Methods

The four methods: Fast_Nextera, Flex, Bio_Flex and Fast_Flex are deribed below after the generic methods. All follow the same principles and can be incorporated into a standard WGS workflow, the differences are in a target enrichment (biotin-streptavidin affinity capture) using methodology from (13) and the number of washes performed during preparation. The advantages to us include cost reduction, improved turnaround times and workflow compatibility leading to increased productivity

### Bacterial Strains and growth conditions

There were two bacterial strains for this work. Firstly, *Escherichia coli* MFD*pir* was used as a plasmid donor strain and cultivated on LB media with the addition of Diaminopimoleic Acid (DAP) to overcome auxotropy (14). Plasmid containing cells were selected for using 100 μg/mL Carbenicillin Disodium (Formedium Ltd. CAR0005) and 50 μg/mL Kannamycin Monosulphate (Formedium Ltd. KAN0005). The recipient strain used was *Escherichia coli* BW25113 (NCTC 14364), this was grown on LB medium with the addition of 50 μg/mL Kannamycin Monosulphate to select for transformants with successful transposition events. For both strains, incubation was at 37 °C.

### Transposon Plasmid Conjugation

Libraries were made using either a Tn*5* family transposon or a mariner family transposon, Himar1. The Tn*5* based libraries were prepared using the conjugative plasmid pBAMD1-2; pBAMD1-2 was a gift from Víctor de Lorenzo (Addgene plasmid # 61564) (15). Similarly, the mariner-based libraries were prepared using the conjugative plasmid pSAM_Ec, pSAM_Ec was a gift from Matthew Mulvey (Addgene plasmid # 10293) (16). Conjugation was performed using a protocol with elements from previously published work (17, 18).The donor and recipient cells were grown overnight to stationary phase in LB broth, with addition of the appropriate supplements or antibiotics. The cells were washed three times in the starting volume of Phosphate Buffered Saline (PBS) to remove any antibiotics and resuspended in 1/20th of the initial volume in PBS. The concentrated donor and recipient cells were then mixed in a ratio of 1:1 volume. The mix was spotted as 10 μL independent matings on an LB agar plate for standard library preparation. The mating plates were incubated at 37 °C for five hours to allow conjugation and transposition to occur.

### Transposon Library Production

Following conjugation, the mating spots were harvested from the LB agar conjugation plates into sterile PBS. The cells were washed twice with PBS and resuspended in 3 mL of PBS. Then, 500 μL of the cell suspension was spread onto a large square bioassay plate (245 mm x 245 mm) of LB agar with 50 μg/mL Kanamycin Monosulphate to select for the transposon containing cells. Six plates were spread per library, aiming for colony density of around 80,000 mutants per plate. The plates were incubated at 37 °C overnight. Following incubation, the plates were harvested from the plates using sterile PBS and a spreader, the mutants from all six plates were pooled to generate one library. An equal volume of 40% (v/v) glycerol was added to give a final glycerol concentration of around 20%, before being stored at -80° C. For this work, nine libraries per transposon were made as described above, 3 biological and technical repeats. The nine libraries were pooled at an equal estimated mutant density to generate the final library pool. Then 10^9^ cells from the final pools were inoculated into 10 mL LB broth, incubated for 18 hours at 37C at 200 rpm, until the cultures reached stationary phase.

### DNA Extraction

For each transposon, DNA was extracted in triplicate from roughly 10^9^ cells using the Maxwell® RSC Cultured Cells DNA extraction Kit (Promega Catalogue No. AS1620) and automated program. The DNA was pooled per transposon, quantified and normalised to 25 ng/uL with molecular grade water and 25 ng was used for each method of sequencing library production.

### Sequencing Primer Design

The custom sequencing primers used in this work were designed with the same rationale as described (19) including the following 5 distinct regions, which are required to be compatible with the Illumina sequencing platform. From 5’ the distinct regions are: the Illumina i5 adapter sequence (29 bp), the Illumina Nextera i5 index (8 bp), the Illumina Read 1 Sequencing primer site (35 bp), Optional Diversity Spacer (8bp), Transposon Binding Site (varying bp).

### Biotinylated Primer Design

A biotinylated primer was designed for use during amplification of the Transposon-Chromosome (Tn-Chr) junctions to enrich and enhance recovery of true junctions and get a greater sequencing depth(13). The primer was designed to be used with the Illumina DNA Prep kit (Illumina 20060060). As such, the sequence was the same as the annealing sequence of the Illumina i7 barcoding primer (20) but with a 5’ biotin added. Sequence: [BTN] –GTCTCGTGGGCTCGG.

### New library preparation methods under evaluation

### Nextera Sequencing Method (FAST_Nextera)

12.5 μl of TD Tagment DNA Buffer (Illumina Catalogue No. 20034198) was mixed with 2.5 μL TDE1, Tagment DNA Enzyme (Illumina Catalogue No. 20034198) and 14 μL PCR grade water was combined to make a tagmentation mix. Genomic DNA was normalised to 25 ng/ μL. 1 μL of normalised DNA (25ng total) was pipette mixed with the 24 μL of the tagmentation mix and heated to 55 °C for 10 minutes in a PCR block.

Then 21 μL kapa2G Fast HS Master Mix (Merck Catalogue No. KK5601) per sample was added with 2 μL of each P7 of Nextera XT Index Kit v2 index primers (Illumina Catalogue No. FC-131-2001 to 2004) and 2 μL of each custom P5 transposon primer (10 mM). Finally, the 25 μL of tagmented DNA was added and mixed. The PCR was run with 72°C for 3 minutes, 95°C for 3 minutes, 28 cycles of 95°C for 10s, 55°C for 20s and 72°C for 1 minute and finally held at 10 °C until ready to proceed.

### Flex Sequencing Method (Flex)

DNA was normalised to 25 ng in 15 μL of molecular grade water. The Illumina DNA Prep kit (Illumina Catalogue No. 20060060) and protocol (21) were used but reactions were minimised to ¼ original volumes; 2.5 μL of BLT and 2.5 μL of TB1 were added to the normalised DNA and mixed. The reaction was incubated in a thermocycler at 55 °C for 15 minutes with the lid set to 100 °C, then cooled to 10 °C. Tagmentation was stopped with the addition of 2.5 μL of TSB, mixed well, and incubated for 15 minutes at 37 °C in a thermocycler with the lid set to 100 °C, then 10 °C. The reactions were incubated on a magnetic plate for five minutes; the supernatant was removed. The beads were then washed twice in 100 μL of TWB and the supernatant removed.

The beads were resuspended in a mix of 20 μL of KAPA2G Fast HS Master Mix, 2 μL of custom P5 transposon primer (10 mM), 2 μL P7 Nextera XT Index Kit v2 index primers The PCR program was 72 °C for 3 minutes, 95 °C for 3 minutes, then 28 cycles of 98 °C for 10 seconds, 55 °C for 20 seconds and 72 °C for 1 minute and finally held at 10 °C until ready to proceed.

### Biotinylation Method (Bio_Flex)

DNA was normalised to 25 ng in 15 μL of molecular grade water. The Illumina DNA Prep kit and protocol (21) were used but reactions minimised to ¼ original volumes ; 2.5 μL of BLT and 2.5 μL of TB1 were added to the normalised DNA and mixed. The reaction was incubated in a thermocycler at 55 °C for 15 minutes with the lid set to 100 °C, then cooled to 10 °C. Tagmentation was stopped with the addition of 2.5 μL of TSB, mixed well, and incubated for 15 minutes at 37 °C in a thermocycler with the lid set to 100 °C, then 10 °C. The reactions were incubated on a magnetic plate for five minutes; the supernatant was removed. The beads were then washed twice in 100 μL of TWB and the supernatant removed.

The beads were resuspended in a mix of of 20 μL of KAPA2G Fast HS Master Mix, 2 μL of Custom P5 transposon primer (10 mM), 2 μL of Biotinylated i7 primer (10 mM.) The PCR program was 72 °C for 3 minutes followed by 16 cycles of 95 °C for 10 seconds, 55 C for 20 seconds and 72 °C for 1 minute and finally held at 4 °C until ready to proceed. The PCR products were purified with a 1:1 ratio of KAPA Pure beads. The final product was eluted in 40 μL of elution buffer (EB) (Qiagen, Hilden, Germany).

The biotinylated products were captured using the DynaBeads Kilobase BINDER kit (Fisher Scientific Catalogue No. 10662244). Using a magnet, 10 μL (100 μg) of Dynabeads were pelleted and the supernatant removed. The beads were washed by resuspending in 40 μL Binding Solution, returned to the magnet, supernatant removed and then resuspended in a further 40 μL of Binding Solution. The Dynabeads were then added to the 40 μL of PCR product from above and incubated on a roller for 4 hours at room temperature; with periodic checks that the beads remained in suspension.

Following incubation, the beads were washes twice with 40 μL Washing Solution and twice with 40 μL distilled water. The beads were finally resuspended in 20 μL EB. Then combined with 25 μL of KAPA2G Fast HS Master Mix, 2.5 μL of Custom P5 transposon primer (10 mM) and 2.5 μL P7 Nextera XT Index Kit v2 index primers. The PCR program was 72 °C for 3 minutes followed by 12 cycles of 95 °C for 10 seconds, 55 °C for 20 seconds and 72 °C for 1 minute and finally held at 10 °C until ready to proceed. The reactions were placed on a magnet and the supernatant containing the amplified target sequences was removed.

### Fast Flex Method (Fast_Flex)

DNA was normalised to 25 ng in 15 μL of molecular grade water. The Illumina DNA Prep kit () and protocol (21) were used but reactions were halved. 2.5 μL of BLT and 2.5 μL of TB1 were added to the normalised DNA and mixed. The reaction was incubated in a thermocycler at 55 °C for 15 minutes with the lid set to 100 °C, then cooled to 10 °C. The reactions were placed on a magnet and the supernatant removed.

The beads were resuspended in a mix of 20 μL of KAPA HiFi 2x PCR Mastermix, 2 μL of Custom P5 transposon primer (10 mM), 2 μL P7 Nextera XT Index Kit v2 index primers. The PCR program was 72 °C for 3 minutes followed by 16 cycles of 98 °C for 10 seconds, 64 °C for 60 seconds and 72 °C for 20 seconds and finally held at 4 °C until ready to proceed.

### Sequencing

The PCR amplification products for all methods were quantified using Qubit High Sensitivity double stranded DNA kit (Catalogue No. 10164582) and run on a Qubit 3.0 instrument and Libraries were pooled following quantification in equal quantities. The pool was double-SPRI size selected between 0.5 and 0.7X bead volumes using KAPA Pure Beads (Roche Catalogue No. 07983298001).

The final pool to be sequences was quantified on a Qubit 3.0 instrument using the High Sensitivity assay kit and run on a D5000 ScreenTape (Agilent Catalogue No. 5067-5588 & 5067-5589) using the Agilent Tapestation 4200 to calculate the final library pool molarity. The prepared TraDIS sequencing libraries were pooled with 937 WGS and 48 PCR amplicon samples to generate a final loading pool for sequencing. The final pool was run on the Illumina NextSeq 2000 instrument on a P4 X-leap 300 cycle kit PE151.

### Analysis

The raw sequence reads were quality trimmed (Q20) and Nextera Transposase adapter contamination was removed using Trim Galore (Version 0.6.7) (22). All of the reads were trimmed using Cutadapt (version 4.9) (23) using the respective sequences 10 bp upstream of the transposon tag. The trimmed reads were then analysed with the Biotradis pipeline (version 1.4.5) (24). Transposon insertions were identified by the tags (TATAAGAGACAG) and (TCCAACCTGT) for Tn*5* and mariner respectively. The reads were mapped to the reference *E*.*coli* BW25113 reference sequence (GenBank: CP009273.1) using smalt within the Biotradis pipeline by calling smalt and the parameters were set as follows; m 1, mapping quality 0, smalt_k 13, smalt_s 1, smalt_y 0.9, smalt_r 0. The insert plot files were then filtered to remove any low level WGS contaminating reads, these were determined as a mapping count of less than 5% of the mean number of reads mapped across all sites per library. Gene essentiality was determined using the BioTradis Gene essentiality command, using R (version 4.2) with the reference genome as GenBank: CP009273.1.

## Results

The four library preparation methods were evaluated for cost, hands-on time in the laboratory, target sequencing and target mapping.

### Cost and workflow comparison

The first step in assessing method selection was to compare the cost of reagents for each methodology. The cost per library varied substantially from £8.81 to £52.38 as seen in **Table 1** with the enzyme format making the significant difference in cost. The Nextera based method was much more expensive with the cost difference being attributed to TDE1 enzyme (previously part of Nextera XT kits). Inclusion of a biotin-streptavidin capture step increased reagent costs by approximately 50% relative to standard Flex protocols, although the absolute increase (∼£5) was modest. **Table 2** breaks down the cost for each method of library production. This was calculated for reagents used specifically in the protocol so excludes plasticware and general laboratory consumables such as nuclease free water and ethanol.

**Table 1.**
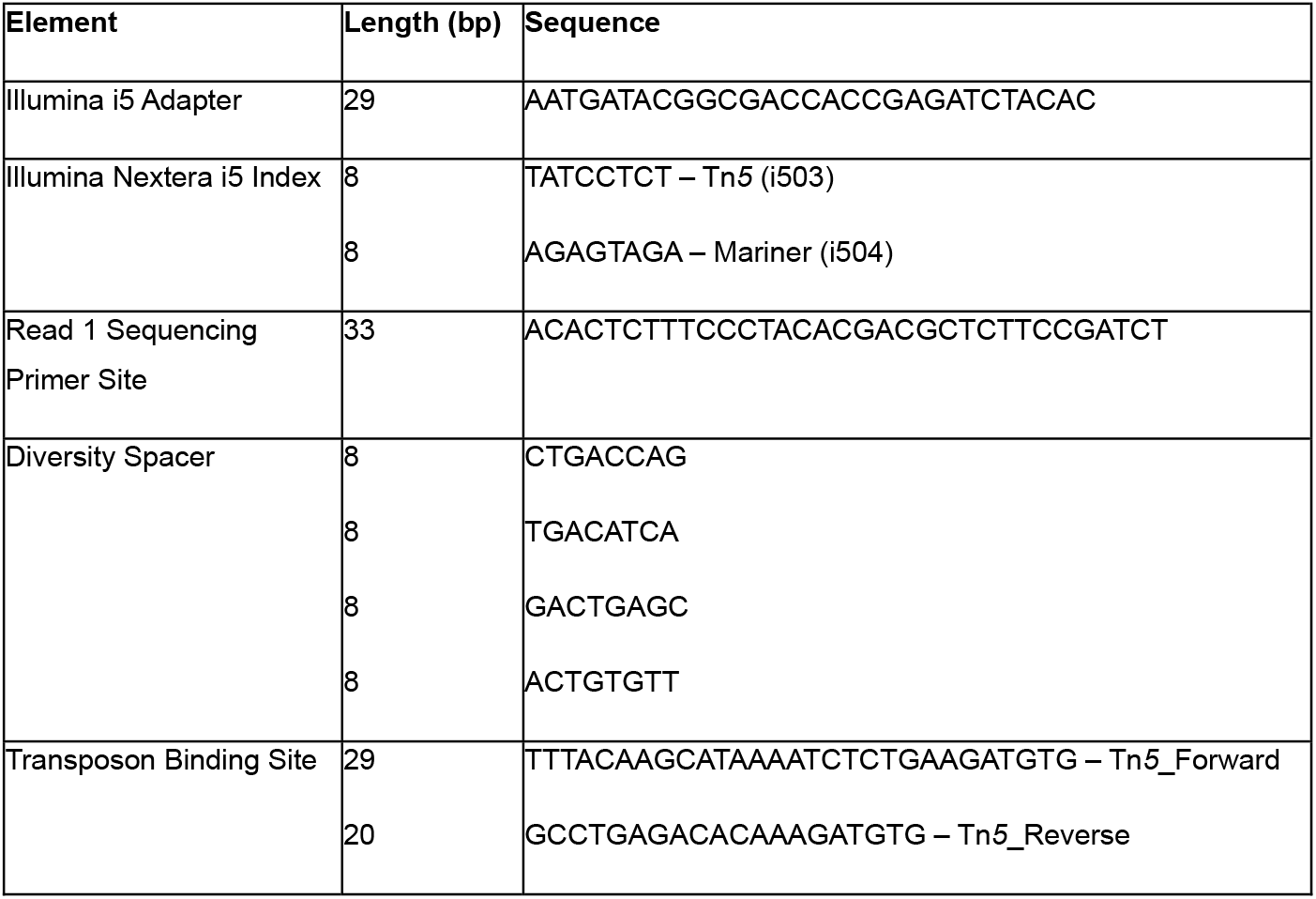
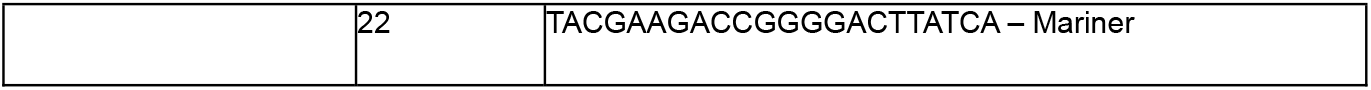
The five elements required for designing a transposon mutagenesis library custom sequencing primer, designed to be used with the Illumina Nextseq platform. The i5 adapter and Transposon binding site are variable regions, with the binding site being specific to the transposon used for the study.

**Table 2:** Cost comparison for four methods. The cost of reagents for one reaction of each preparation method, costs are in GDP and correct as of April 2025. (-) indicates reagents not required for the specified protocol.

| Reagent | Fast_Nextera | Flex | Bio_Flex | Fast_Flex |
| --- | --- | --- | --- | --- |
| DNA Prep Kit * | 48.68 | 7.45 | 7.45 | 7.45 |
| Kapa 2G Master Mix | 0.90 | 0.53 | 1.60 | 0.53 |
| Index primer | 0.01 | 0.01 | 0.01 | 0.01 |
| Tn primer | 0.01 | 0.01 | 0.02 | 0.01 |
| Biotinylated primer | - | - | 0.02 | - |
| Qubit | 0.81 | 0.81 | 0.81 | 0.81 |
| Kilobase Binder Kit | - | - | 3.52 | - |
| Kapa Mag Beads | 1.98 | - | - | - |
| **Total** | **52.57** | **8.81** | **13.43** | **8.81** |

\*TDE1 Enzyme purchased for Fast\_Nextera method, for all Flex based approaches the standard Illumina DNA Prep Kit is used.

The next step was to compare hand-on time. As the DNA preparation and sequencing on the machine were the same for all methods, we looked at the time from normalised genomic DNA to a pooled library ready to be sequenced.

Hands on time and total protocol duration also showed marked differences for some of the protocols, visualised in **Figure 1** and detailed in **Table 3**. Fast_Nextera and Fast_Flex required approximately 2.5 hours from input DNA to sequencing; followed by the standard flex method taking around 3.5 hours. However, the Bio_Flex method spanned 2 working days and doubled the hands-on time compared to the Fast_ methods.

**Table 3:**
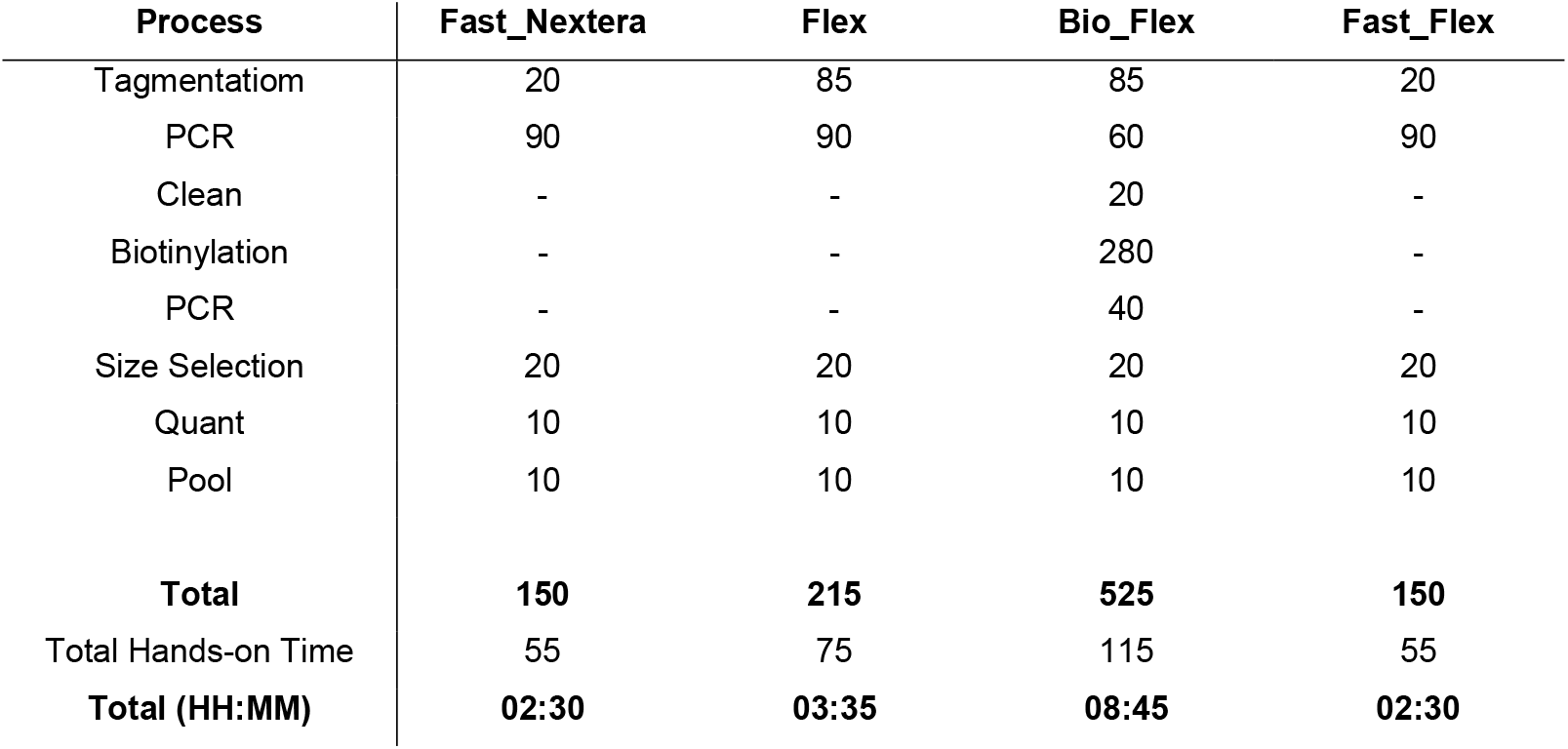
Hands on Time comparison for four methods, The time from pure normalised genomic DNA to final pool prior to sequencing. The values are reported as minutes unless otherwise stated. (-) indicates procedures not required for the specified protocol.

| Process | Fast_Nextera | Flex | Bio_Flex | Fast_Flex |
| --- | --- | --- | --- | --- |
| Tagmentation | 20 | 85 | 85 | 20 |
| PCR | 90 | 90 | 60 | 90 |
| Clean | - | - | 20 | - |
| Biotinylation | - | - | 280 | - |
| PCR | - | - | 40 | - |
| Size Selection | 20 | 20 | 20 | 20 |
| Quant | 10 | 10 | 10 | 10 |
| Pool | 10 | 10 | 10 | 10 |
| **Total** | **150** | **215** | **525** | **150** |
| Total Hands-on Time | 55 | 75 | 115 | 55 |
| **Total (HH:MM)** | **02:30** | **03:35** | **08:45** | **02:30** |

**Figure 1.**
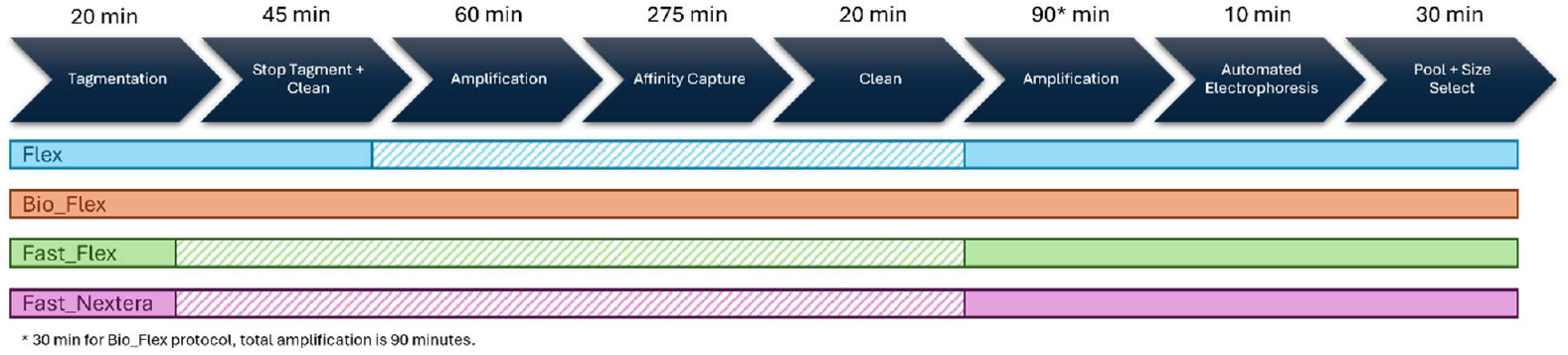
A graphical overview of the four methods being compared and where the differences in the processes are.

### Target enrichment and mapping performance

The four methods were all successful in generating TIS sequencing libraries across both of the transposons used, the BioTradis output is summarise in **Table 4**.

**Table 4.** Target sequencing and mapping comparison for four methods. BioTraDIS Output table

|  | Fast_Nextera | Flex | Bio_Flex | Fast_Flex |
| --- | --- | --- | --- | --- |
| **Tn5** |  |  |  |  |
| Total reads | 2.32e+07 | 3.91e+07 | 3.29e+07 | 3.70e+07 |
| % Matched | 67.56 | 78.63 | 94.96 | 79.79 |
| % Mapped | 58.51 | 46.89 | 62.25 | 45.19 |
| Unique insertion sites | 1.43e+05 | 1.78e+05 | 1.89e+05 | 1.21e+05 |
| Average distance (bp) | 32.31 | 26.01 | 24.54 | 38.29 |
| Filtered insertion sites | 7.31e+04 | 8.29e+04 | 7.62e+04 | 6.62e+04 |
| Filtered average distance (bp) | 63.40 | 55.86 | 60.80 | 69.93 |
| **Mariner** |  |  |  |  |
| Total reads | 3.44e+07 | 4.09e+07 | 6.35e+07 | 2.89e+07 |
| % Matched | 87.01 | 84.67 | 96.55 | 82.94 |
| % Mapped | 44.27 | 37.96 | 55.04 | 38.51 |
| Unique insertion sites | 3.28e+05 | 2.29e+05 | 3.02e+05 | 2.07e+05 |
| Average distance (bp) | 14.10 | 20.21 | 15.32 | 22.41 |
| Filtered insertion sites | 2.41e+05 | 1.74e+05 | 1.87e+05 | 1.53e+05 |
| Filtered average distance (bp) | 19.20 | 26.66 | 24.83 | 30.33 |

The Flex based protocols outperformed the Nextera method for enrichment of Tn-Chr junctions in the Tn*5* preparations and performed, measured by the percentage of reads matched in **Table 4**. This was not the case for the mariner preparations, but the differences were minimal (4% mapping) compared to the Tn*5* (11% mapping) from Nextera and the lowest mapping flex. Biotinylating did achieve the highest enrichment across both the Tn*5* and mariner libraries, although this did not consistently translate into a higher number of filtered unique insertion sites.

Biotinylation and Illumina’s flex protocols provided the beset Tn-Chr junction enrichment determined by the percent matched. Bio_Flex and Flex performed similarly for the Tn*5* library but were the best methods when judged by unique insertion counts. For the mariner library, Bio_Flex and Nextera were the best when judged by unique insertion count. Across both transposons, the Nextera method was the least efficient relative to cost, **Table 2**, producing lower enrichment and fewer informative reads.

### Sequencing Depth Required

The major cost of sequencing a transposon comes from data generation, the more data required, the more cost. As seen in **Table 4**, more sequence reads do not guarantee more unique insertions. Using the sequences generated for the comparisons above, we concatenated and then randomly subsampled the sequences to simulate depths of 1M reads to 100M reads. The subsampling was performed in triplicate. The cost per Gb of data was estimated to around £5 per million reads.

As expected, more sequence reads identify more unique insertions. However, subsampling showed diminished returns, particularly with the unfiltered data, darker colours of **Figure 2**. Especially for Tn*5*, an increased number of reads led to a greater number of unique insertions but gave fewer essential genes and therefore less biologically relevant information. The greater spread of insertion events across the genome leads to fewer statistically significant insertions withing coding sequences in the genome and therefore the BioTradis package does not deem these sequences to be essential. When filtering, low level WGS reads were bioinformatically removed. The effect of filtering was far less pronounced for the mariner libraries; this is because mariner inverted repeats have no overlap with the Tn*5* mosaic ends used by Illumina. The overlap is at the 3’ end of the sequencing primer for Tn-Chr enrichment, **Table 1**, has no overlap with the Illumina adapter sequences. Previous works (25-28) report that there are 300-700 essential genes in *E*.*coli* BW5113 or related MG1655 (29).

**Figure 2.**
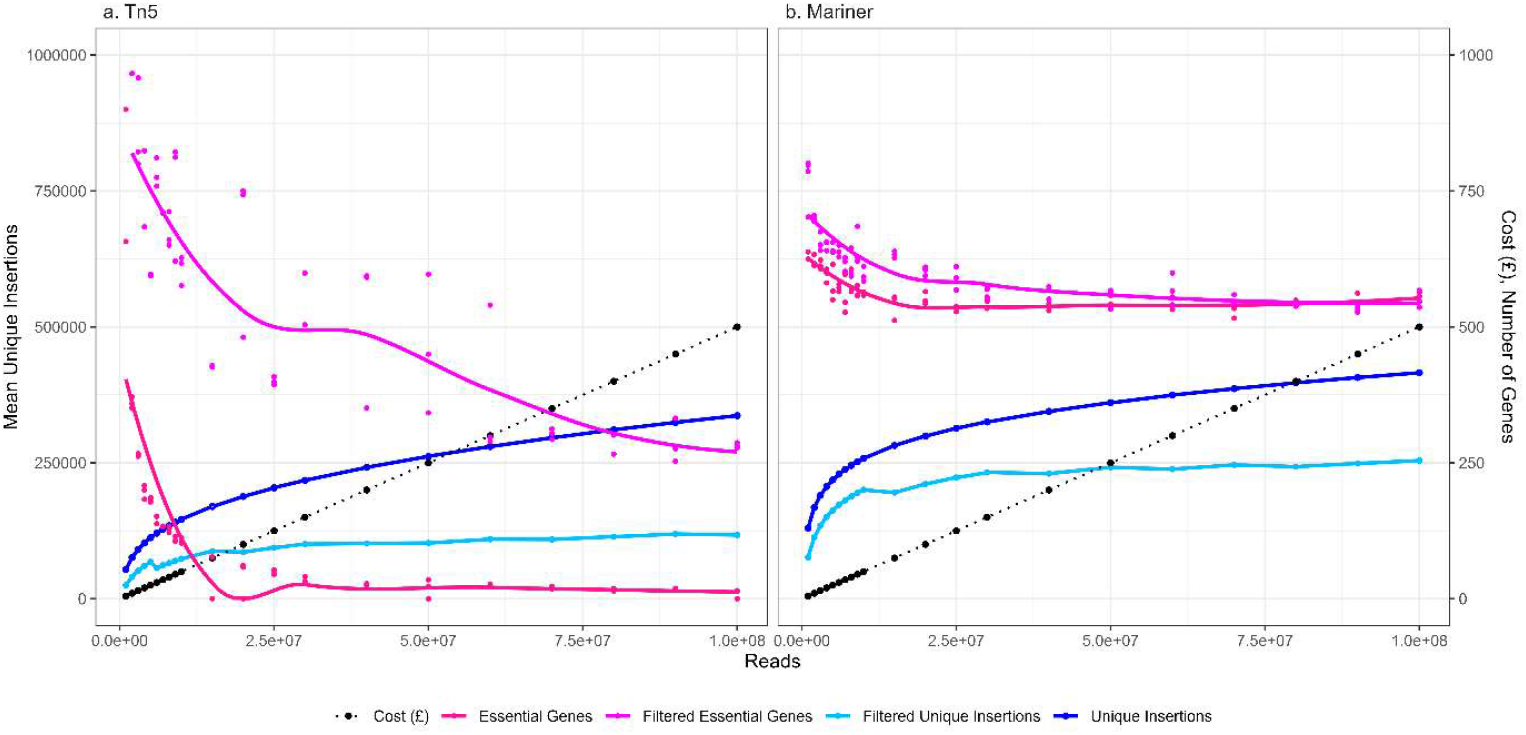
Simulated sequence depth analysis for unique insertion, essential genes and costs of **(a)** Tn5 and **(b)** mariner libraries used in this work, sequenced at depths from 1 million to 100 million reads showing the cost of data at each depth, the number of unique insertions and essential genes for the unfiltered (dark) and filtered (light) plot files.

The analyses of libraries used in this work indicate that a sequence depth of around five million reads is sufficient to statistically identify essential genes, especially for the insertion density of the mariner library with a cost of £25 for data generation. Beyond this depth, there was no biological insight gained form deeper sequencing . For Tn*5*, the libraries have far fewer unique insertions, therefore a greater sequencing depth was required to capture the true insertions and eliminate the background noise by filtering. With the Tn*5* sequencing depths **Figure 2a**, it was evident that form around 25M to 50M, £125 - £250 reads was the optimal depth for sequencing before the number of spurious background reads overwhelmed the statistical analyses.

## Discussion

This study demonstrates that simplified Illumina based sequence library preparation methods generate robust TIS data with Flex based protocols being a low-cost compared to traditional Nextera based protocols. All four approaches generated similar data with the majority of variation being the sequencing depth and the number of mutants within the TIS library.

We have tested the protocols with two routinely used transposon families, Tn*5* and mariner, the protocols are more efficient with mariner than Tn*5*, **Figure 2**, if using a universal method. However, improvements can be made in the enrichment step following Tagmentation by optimising primer design, annealing temperatures and extensions times as you would a normal PCR reaction. While they were the only two transposons tested, the methods used here will be applicable to any family of transposon with appropriate primer design. The primers in this study, **Table 1**, have the inclusion of a diversity spacer, incorporating an element of random built in which removes the need for dark cycles (7, 24) and overcomes a common problem seen with amplicon sequencing, particularly TIS. Since WGS has natural diversity, this compensates for the lack in TIS, so this element is no longer a requirement, this work used primers designed prior to this work.

When considering the cost of sequence preparation alone, Nextera-based preparation costs 4 or 6 times compared to the Flex approaches. Nextera preparation of sequencing libraries for TIS is commonplace and generally the approach used, there are other Tagmentation methods that follow a similar laboratory workflow but use a Mu family transposon for the Tagmentation procedure(13, 30); while this is comparable in cost and time for the Flex protocols, the product has been discontinued by the manufacturer (Thermo Fisher). While other commercial kits are available for Illumina sequencing library production, these were also not included in this study as there was sufficient evidence that the Illumina reagents were producing good results. The Nextera based protocol has been used historically (19, 31), as the workflow was complimentary to Illumina. However, in 2018 Illumina changed to Flex protocols, the Nextera approach is still compatible but uses outdated prep kits or requires the costly TDE1 enzyme.

For the three flex protocols, the added cost of biotinylation is an extra 50% compared to the other two methods, **Table 2**, but this represents around £5 so is negligible when considering the overall cost of an experiment. The negating factor for the biotinylation protocol is that it takes double the time of the next longest protocol (Flex). This is not reflected in the hands-on time required but this too is increased for the biotinylation protocol. While data presented here shows that biotinylating performed best at Tn-Chr enrichment (highest percentage of reads matching the transposon tag) it did not always provide the most unique insertions.

Due to the time required and extra steps involved, the biotinylation approach may not be the most applicable for most TIS sequencing library preparations. Increasing manipulations increases the chance to introduce biases and errors in preparations. Where biotinylating would prove to be particularly useful would be when doing multi organism experiments or where the transposon DNA is likely to be a very small proportion of the DNA. For example, studying survival of mutants in bacterial communities (32) or where there is potentially animal (33) or plant contaminating DNA (34).

Other work, using an adapter ligation approach but with minimised costs demonstrates that decreasing the sequencing costs of hight throughput genomic screening experiments a universal goal (35). However, adaptor ligation approaches have increased complexity and an increase in the number of manipulations compared to tagmentation approaches. This makes tagmentaion preferable and more robust for genomic coverage than adapter ligation (36).

Using a method that is lower in costs means that more replicates can be performed for the same overall cost, providing more robust results.

An important factor to consider when designing a TIS sequencing experiment is the depth of sequencing required. The majority of the cost associated is data generation, with our working estimate being £5 per Gigabyte of data. For a whole genome sequence, the desired coverage, genome size and read length can be used to determine the desired output (37).

For TIS, this calculation is based on estimated library density and a minimum coverage of those mutants; making the calculation is dependent on the individual mutant libraries and experimental set up. The Tn*5* library used for this work would be prohibitively expensive to sequence for use for experimental work, in an experimental set up, more mutants would be made to increase the number of unique insertion sites. In this case, **Figure 2a** would more resemble **Figure 2b** where the reads from true insertions dilute out the noise.

Through this work, we found that the sequencing depth had an impact on essential gene determination. The reporting of essential genes is dependent on sequence depth and with an emphasis on cost, five million reads produced sufficient data, especially in the case of mariner, to determine essential genes. Moreover, increased sequencing depth overwhelmed the statistical analyses leading to genes being misclassified as non-essential. This was most likely attributed to off target primer annealing due to the experimental transposon and Illumina sequencing both using Tn*5*.

While gene essentiality is an active area of research, most TIS libraries used are investigating conditional essentiality under differing stresses or phenotypes (30, 33, 38-40). This work has not explored the impact of sequence library preparation on comparative analysis, however the filtering and mapping, **Table 4** is the first stage of this analysis indicating that any of the four preparation approaches used in this work would yield suitable data for comparative analyses.

## Conclusion

The Illumina based Flex protocol outlined in this work offers a significant cost reduction in the preparation of sequencing libraries for large scale TIS. Two Flex approaches were tested, both cost the same amount and performed well in this work, the standard approach performed marginally better in terms of Tn-Chr junction enrichment and unique insertion count but takes an extra hour of preparation time and involves an extra incubation and wash step. While including an affinity capture step (biotinylation) in the protocol did enhance unique insertion recovery, the 50% extra cost for library preparation (£5) and increased, more than double, preparation time did not offer any real benefit to the number of unique insertions identified. The most influential factor on the number of unique insertions recovered was the depth of sequencing performed. However, more sequence reads, and more unique insertions do not necessarily equate to more biologically relevant conclusions for example gene essentiality. Using the methods and sequencing depth described here would enable TIS of a dense transposon mutagenesis library for a cost of around £40 per sample.

## Data Availability

The reads representing the raw sequence data and the subsampling data sets used in this project can be accessed at **10.5281/zenodo.18339183**

Subsampled reads and scripts used for the TIS analyses, filtering and subsampling are available upon request from Claire Hill

## Acknowledgement of Funding

The author(s) gratefully acknowledge the support of the Biotechnology and Biological Sciences Research Council (BBSRC); this research was funded by BBSRC Institute Strategic Programme Grant Microbes in the Food Chain BB/R012504/1 and the BBSRC Institute Strategic Programme Microbes and Food Safety BB/X011011/1 and its partner project project(s). CH gratefully acknowledges the Medical Research Council (MRC) funding of PhD project MR/R015937/1.

